# Genome-wide association study of alcohol consumption in Russian population

**DOI:** 10.1101/498824

**Authors:** Igor Nizamutdinov, Yaroslav Popov, Dmitriy Romanov, Ekaterina Surkova, Valery Ilinsky, Alexander Rakitko

## Abstract

Liability to alcohol dependence is heritable, but little is known about its complex polygenic architecture. We performed a genome-wide association study (GWAS) of self-reported alcohol consumption in 1798 of Russian individuals. No SNP reached genome-wide significance for any alcohol drinking patterns: never drinking, everyday drinking, once per week drinking and once per month drinking. Polymorphisms in previously reported genes of *KLB* and *AUTS2* were significant associated with everyday drinking and once per week drinking patterns, respectively. We also found associations of genes involved in nervous system function and mental disorders with some alcohol drinking patterns. This study identifies novel gene associations that should be the focus of future studies investigating the neurobiology of alcohol consumption.

## Introduction

Alcohol consumption (AC) in Russia was 13.5 L per capita in 2016. Although AC decreased in Russia previous years [1], it is still one of the major cause of death [2]. Over 200 diseases are linked to AC including stroke [3], cardiovascular events [4] and some cancer types [5].

There is a substantial genetic component to the variation in AC. A recent twin study estimated the heritability of AC to be 0.43 (95% confidence interval 0.31–0.56). In a sample of unrelated parents, ~18% of the variance in AC was attributed to common SNPs in the same study [6].

Specific genetic variants are linked to variation in AC, the most important of them are rs671 in the aldehyde dehydrogenase (*ALDH2*) gene and a cluster of variants spanning the alcohol dehydrogenase genes (*ADH1B, ADH1C, ADH5, ADH6, ADH7*). SNP rs671 leads to slower metabolism of acetaldehyde that causes the alcohol flush reaction [7]. This polymorphism is highly protective against alcoholism in Asian populations [8]. The rs1229984 in *ADH1B* is also associated with an alcohol-flush reaction in Asian populations and thus protects against high alcohol consumption [9]. It is also found to be associated with drinking pattern phenotypes in African and European populations [10].

Previous genome-wide association studies (GWAS) of AC have also found few replicable loci outside of the alcohol metabolizing genes. The rs6943555 in *AUTS2* was shown to be associated with AC [11]. Although it was not replicated in more recent study [12]. A meta-analysis included >105,000 individuals of European ancestry revealed that rs11940694 in *KLB* gene is associated with alcohol intake [12].

Genetic factors of AC vary by race and ethnicity [13]; thereby GWAS in different population are needed to reveal the genetic pattern of AC.

To identify loci associated with AC and replicate an association of previously reported polymorphisms in Russian population, a genome-wide association study was performed.

## Methods

### 1. Subjects

Participants were drawn from the DNA biobank of Genotek Ltd., a direct-to-consumer genetics company. All samples were genotyped using customized Illumina InfiniumCoreExome 24 v1.1 beadchip with 550,000 standard plus 12,000 add-on SNPs or Illumina Global Screening Array v.1 and v.2. After excluding participants with age < 25 years old there were 1798 samples with an average age of 40.2 ± 11.05. The group includes 42% female samples. All participants filled out the online survey with the question “How often do you drink alcohol?” with available answers “Never”, “Every day”, “Once per week” and “Once per month”. The survey was carried out online and all ethical requirements were covered by providing with informed consent of participants. Data used in our analysis were collected prior to December, 2018.

### 2. Data analysis

After genotyping we excluded samples with call rate < 0.1 and SNP with call rate < 0.05. We calculated allele frequencies in the combined sample set and excluded SNPs with minor allele frequency < 0.05 thereafter. Using PRIMUS we determined relatives in our samples and excluded all samples with the relationship degree < 3 except founder. Also we excluded all samples with self-reported non-European ancestry. After all exclusions there were 1008 samples. Using plink, p-value and 95% confidence intervals for the OR (odds ratio) were obtained in the association test.

## Results and Discussion

The distributions of the AC in our sample set are presented in Table 1.

**Table 1.** The distribution of AC in our samples.

| Alcohol drinking patterns | % (N) |
| --- | --- |
| Never | 19.2% (212) |
| Everyday drinking (EDD) | 8.2% (90) |
| Once per week drinking (OWD) | 35.4% (391) |
| Once per month drinking (OMD) | 37.2% (411) |

First, we compare allele frequencies between groups of never drinking and EDD. No SNP gained genome-wide significance. Top associated SNPs are present in Table 2.

**Table 2.** Results for the top loci associated with EDD vs. never drinking status

| SNP | Gene | Alleles | MAF EDD | MAF never drinking | OR [C.I.] | P value |
| --- | --- | --- | --- | --- | --- | --- |
| rs1584108 | intergenic | G/A | 0.61 | 0.22 | 5.41 [2.58, 11.35] | 3.1e-6 |
| rs7782542 | intergenic | T/C | 0.25 | 0.10 | 3.13 [1.82, 5.38] | 2.0e-5 |
| rs4798131 | DLGAP1 | C/T | 0.33 | 0.52 | 0.45 [0.31, 0.66] | 2.6e-5 |
| rs714584 | CNTNAP2 | T/G | 0.33 | 0.52 | 0.45 [0.31, 0.66] | 2.6e-5 |
| rs11680427 | ANTXR1 | C/T | 0.65 | 0.29 | 4.49 [2.16, 9.32] | 3.1e-5 |
| rs7550535 | intergenic | T/C | 0.65 | 0.27 | 5 [2.27, 11.02] | 3.4e-5 |

\* Alleles present minor/major, OR calculated for minor allele

The rs4798131 in *DLGAP1* was associated with everyday drinking in our group. *DLGAP1* encodes SAPAP1 protein localized at postsynaptic density and involved in maintaining normal brain function and development [16]. Previously polymorphisms in *DLGAP1* were not shown to be associated with alcohol behaviour, however they are associated with schizophrenia and other brain diseases [17].

We found the association rs714584 in *CNTNAP2* gene with everyday drinking comparing with never drinking group. Recently some SNPs in *CNTNAP2* were shown to be associated with alcohol addiction in female [18]. This gene functions in nervous system as cell adhesion molecules and receptors. It also associated with epilepsy [19], autistic disorder [20] and major depression disorder [21].

The SNP rs3102165 in *KLB* gene was associated with everyday drinking with p = 4.9e-3. Also this SNP is not in linkage disequilibrium (LD) with rs11940694 (R^2^ = 0.008).

Second, allele frequencies were compared between OWD and never drinking groups. The top results are present in Table 3.

**Table 3.** Results for the top loci associated with OWD vs. never drinking status

| SNP | Gene | Alleles* | MAF OWD | MAF never drinking | OR [C.I.] | P value |
| --- | --- | --- | --- | --- | --- | --- |
| rs7781928 | STEAP2-AS1 | A/G | 0.42 | 0.27 | 1.96 [1.49, 2.57] | 1.3e-6 |
| rs6088792 | UQCC1 | T/C | 0.22 | 0.46 | 0.33 [0.20, 0.55] | 1.7e-5 |
| rs13428332 | intergenic | G/A | 0.07 | 0.15 | 0.45 [0.31, 0.65] | 2.3e-5 |
| rs75615630 | PRKCA | C/T | 0.22 | 0.11 | 2.43 [1.59, 3.69] | 2.4e-5 |
| rs35565362 | intergenic | A/G | 0.42 | 0.27 | 1.94 [1.42, 2.64] | 2.4e-5 |
| rs2890956 | intergenic | G/A | 0.36 | 0.51 | 0.54 [0.41, 0.72] | 2.8e-5 |

\* Alleles present minor/major, OR calculated for minor allele

The polymorphism of rs75615630 in PRKCA was associated with OWD in our samples. Protein kinase C alpha encoded by PRKCA gene is involved in many processes in nervous system including synaptic signalling [22], neurite growth and neuronal development [23], and possibly in myelination [24]. It also plays role in memory formation including working memory [25].

The rs740114 in AUTS2 was associated with OWD in our sample with p = 1.1e-3. This SNP is not in LD with rs6943555 (R2 = 0.001).

Then we compared allele frequencies between OMD and never drinking groups. The top results are present in Table 4.

**Table 4.** Results for the top loci associated with OMD vs. never drinking status

| SNP | Gene | Alleles* | MAF OMD | MAF never drinking | OR [C.I.] | P value |
| --- | --- | --- | --- | --- | --- | --- |
| rs1999923 | intergenic | A/G | 0.25 | 0.43 | 0.43 [0.31, 0.61] | 1.7e-6 |
| rs877818 | EFNB1 | A/G | 0.12 | 0.24 | 0.43 [0.29, 0.62] | 3.9e-6 |
| rs6593376 | LINC0084<br>1 | T/C | 0.13 | 0.25 | 0.45 [0.32, 0.64] | 4.3e-6 |
| rs7880190 | intergenic | C/T | 0.25 | 0.42 | 0.45 [0.32, 0.64] | 4.5e-6 |
| rs17360994 | KDF1 | C/T | 0.07 | 0.15 | 0.43 [0.30, 0.64] | 1.4e-5 |
| rs2334501 | KRTAP5-2 | T/C | 0.18 | 0.44 | 0.27 [0.15, 0.50] | 1.4e-5 |

\* Alleles present minor/major, OR calculated for minor allele

Finally, we tried to discover genetic markers associated with never drinking status. We compare allele frequencies between never drinking group and EDD, OWD and OMD together. The results are in Table 5.

**Table 5.** Results for the top loci associated with never drinking status

| SNP | Gene | Alleles* | MAF never drinking | MAF EDD, OWD and OMD | OR [C.I.] | P value |
| --- | --- | --- | --- | --- | --- | --- |
| rs17360994 | KDF1 | C/T | 0.15 | 0.08 | 2.11 [2.92, 1.52] | 4.5e-6 |
| rs1999923 | intergenic | A/G | 0.43 | 0.27 | 1.99 [2.70, 1.47] | 5.9e-6 |
| rs11854732 | intergenic | C/T | 0.32 | 0.20 | 1.84 [2.42, 1.40] | 8.7e-6 |
| rs1350406 | intergenic | G/A | 0.49 | 0.37 | 1.62 [2.01, 1.31] | 8.8e-6 |
| rs35414786 | TSPAN14 | T/C | 0.29 | 0.17 | 1.91 [2.55, 1.43] | 9.0e-6 |
| rs12742376 | KDF1 | T/C | 0.15 | 0.08 | 2.02 [2.79, 1.47] | 1.3e-5 |

\* Alleles present minor/major, OR calculated for minor allele

We found two linked SNPs (R^2^ = 0.991) in *KDF1* gene associated with never drinking status. This gene was not implicated to AC or alcohol use disorders in previous studies and its role in nervous system is unclear. Some mutation in KDF1 cause hypohidrotic ectodermal dysplasia [26].

## Conclusion

This study presents the GWAS of alcohol consumption in Russian population and identifies several genetic loci in genes involved in nervous system function and mental disorders with some alcohol drinking patterns. Further independent studies are required to confirm these findings.

